# Characterizations of Antibacterial and Anti-biofilm Properties of Dendrimers G6 and G9

**DOI:** 10.64898/2026.06.02.729742

**Authors:** Thushara Galbadage, Gideon Igo, Yuchang Chen, Rafael Nhancale, Katelyn Spradley, Hyun Ho Chung, Richard S. Gunasekera

**Affiliations:** Department of Health Sciences and Public Health, Texas Christian University, 2901 Stadium Drive, Fort Worth, TX 76129, USA; Department of Chemistry and Biochemistry, Biola University, 13800 Biola Ave, La Mirada, CA 90639, USA

**Keywords:** antimicrobial resistance, minimum inhibitory concentration (MIC), cationic dendrimers, nanomedicine, bacterial pathogens, MRSA, *P. aeruginosa*, biofilm inhibition

## Abstract

Dendrimers are nanosized molecules with potential antimicrobial applications. This study evaluates the antibacterial and anti-biofilm properties of two cationic dendrimers, NVX-G6 (G6) and NVX-G9 (G9), against clinically relevant bacterial pathogens. Minimum inhibitory concentrations (MIC) and minimum bactericidal concentrations (MBC_99_) were determined for *Escherichia coli, Pseudomonas aeruginosa*, methicillin-sensitive *Staphylococcus aureus* (MSSA), and methicillin-resistant *Staphylococcus aureus* (MRSA). The synergy of dendrimers with ceftazidime and vancomycin was evaluated using checkerboard assays. Furthermore, biofilm formation inhibition assays and fluorescent microscopy were performed to assess dendrimer interactions with bacterial biofilms. The results indicate that G6 and G9 exhibit limited direct antibacterial activity at high concentrations (MIC > 1024 µg/mL) but demonstrate synergistic effects when combined with ceftazidime against *E. coli* and *P. aeruginosa* (FIC < 0.5). Notably, both dendrimers penetrated and colocalized within established biofilms, with time-dependent reductions in biomass observed after extended incubation, suggesting a role in progressive biofilm disruption rather than acute inhibition of formation, although significant biomass reduction was not observed under standard assay conditions. These findings contribute to the understanding of dendrimer-antibiotic interactions and their implications in antimicrobial and nanomedicine therapy.

## INTRODUCTION

The rise of antimicrobial resistance is a global health crisis that necessitates innovative therapeutic interventions (Prestinaci et al., 2015; Salam et al., 2023; The Lancet Respiratory, 2024). Traditional antibiotics are losing efficacy due to the emergence of resistant bacterial strains, leading to increased mortality and healthcare costs (Ahmed et al., 2024). Among the strategies explored to combat resistance, nanomedicine offers promising alternatives. Dendrimers, hyperbranched macromolecules with tunable chemical properties, have been extensively studied for their potential in drug delivery, gene therapy, and antimicrobial applications (Abbasi et al., 2014; Mittal et al., 2021).

Dendrimers have emerged as a promising class of nanoscale macromolecules because their highly branched structure, multivalent surface groups, and tunable functionalization enable interactions with bacterial cell membranes and extracellular matrices in ways distinct from traditional antibiotics. Several generations of poly(amidoamine) (PAMAM) and poly(propylene imine) dendrimers have demonstrated antimicrobial properties through mechanisms such as membrane permeabilization, charge-based disruption, induction of oxidative stress, and improved antibiotic delivery. Despite these advantages, dendrimer efficacy has varied widely across studies, and higher-generation dendrimers often exhibit increased cytotoxicity, necessitating a deeper understanding of structure and activity relationships when applied to living microbial systems.

Cationic dendrimers, such as NVX-G6 (G6) and NVX-G9 (G9), interact with bacterial membranes through electrostatic forces, potentially leading to membrane disruption and bacterial cell death (Santos et al., 2019). However, their efficacy against different bacterial species and their role in biofilm-associated infections remain underexplored. Biofilms, structured communities of bacteria encased in a self-produced matrix, contribute significantly to chronic infections and antibiotic resistance (Sharma et al., 2019; Shree et al., 2023). Understanding whether dendrimers can inhibit or disrupt biofilms is critical in developing alternative therapeutic approaches.

Biofilm-associated infections represent one of the most treatment-refractory manifestations of antimicrobial resistance. Biofilms on lung tissue, wounds, and indwelling devices such as catheters, prosthetics, and ventilators display antibiotic tolerance up to 1,000-fold higher than planktonic cells, contributing to persistent infections and elevated morbidity. *P. aeruginosa* and *S. aureus*, two of the most clinically significant pathogens studied here, routinely form dense, protective biofilms that limit antibiotic penetration and facilitate horizontal gene transfer. Because of this, there is growing interest in nontraditional agents, such as cationic dendrimers, that may disrupt, penetrate, or sensitize biofilms to existing antibiotics.

While cationic dendrimers have been recognized for their membrane-interactive properties, their functional role in modulating antibiotic susceptibility and biofilm formation remains insufficiently characterized. Very few studies have evaluated dendrimers with limited inherent antimicrobial activity to determine whether they can nonetheless potentiate antibiotic performance or alter biofilm architecture. Additionally, the specific dendrimers examined here, NVX-G6 and NVX-G9, represent two higher-generation, highly cationic structures that may exhibit distinct physicochemical interactions with bacterial membranes and extracellular polymeric substances. However, their antibacterial potency, synergy potential, and capacity to localize within or affect biofilms have not previously been investigated. This lack of data represents a critical barrier to understanding whether dendrimers should be developed primarily as standalone antimicrobials, antibiotic adjuvants, or anti-biofilm agents.

Therefore, this study aims to: (1) characterize the antibacterial activity of G6 and G9 against clinically relevant Gram-negative and Gram-positive pathogens, (2) determine whether these dendrimers enhance antibiotic susceptibility through synergistic interactions with ceftazidime or vancomycin, and (3) evaluate their effects on biofilm formation, biomass, and dendrimer localization within biofilm matrices. By addressing these knowledge gaps, this work advances the understanding of how dendrimer-antibiotic interactions may be leveraged to overcome antimicrobial resistance and improve therapeutic outcomes.

To achieve these objectives, we employed standardized antimicrobial susceptibility testing, checkerboard synergy assays, quantitative biofilm assays, and fluorescence imaging to assess dendrimer-bacteria interactions across multiple biological contexts. The following sections describe the experimental procedures used to evaluate dendrimer activity and their potential as adjunctive antimicrobial or anti-biofilm agents.

## MATERIALS AND METHODS

### Bacterial Strains and Growth Conditions

Bacterial strains used in this study included *Pseudomonas aeruginosa* ATCC 27853, *Escherichia coli* ATCC 11303, methicillin-sensitive *Staphylococcus aureus* (MSSA) ATCC 25923, and methicillin-resistant *Staphylococcus aureus* (MRSA) ATCC 33591. Bacterial cultures were maintained in Luria-Bertani (LB) broth or Tryptic Soy Broth (TSB) and incubated at 37°C under aerobic conditions. For experimental use, overnight bacterial cultures were diluted 1:100 in fresh LB or TSB to obtain secondary cultures with an optical density at 600 nm (OD600) of ∼0.1. These secondary cultures were used for all subsequent antimicrobial and biofilm assays.

### Dendrimer Preparation

Two cationic dendrimers, NVX-G6 (G6) and NVX-G9 (G9), were evaluated for their antibacterial and antibiofilm properties. Stock solutions of G6 and G9 were prepared in sterile deionized water at a concentration of 64 mg/mL and stored at -20°C until use. Dendrimer working solutions were prepared fresh before each experiment by serial dilution in the appropriate media.

### Microdilution Assays

The microdilution assay was performed following Clinical and Laboratory Standards Institute (CLSI) guidelines to determine the minimum inhibitory concentration (MIC) of G6 and G9. Briefly, two-fold serial dilutions of dendrimers were prepared in Mueller-Hinton Broth (MHB) in a 96-well microtiter plate, achieving final concentrations ranging from 2 to 1024 µg/mL. Bacterial suspensions at ∼5 × 10^5^ CFU/mL were added to each well, and plates were incubated at 37°C for 24 h. MIC was defined as the lowest concentration of the dendrimer that inhibited visible bacterial growth. To determine the minimum bactericidal concentration (MBC99), aliquots from wells showing no visible growth were plated onto LB agar, and the lowest concentration that resulted in a 99% reduction in CFU was recorded.

### Disc Diffusion Assays

The antimicrobial activity of G6 and G9 was also evaluated using the disc diffusion assay and compared with similar disk diffusion assays performed using Ampicillin, Gentamycin, and Kanamycin (Figure 1). Sterile filter paper discs (6 mm diameter) were impregnated with 10 µL of dendrimer solutions at concentrations of 64, 128, 256, and 512 µg/mL. The discs were placed onto Mueller-Hinton agar plates inoculated with bacterial suspensions standardized to 0.5 McFarland units. Plates were incubated at 37°C for 24 h, after which the zone of inhibition (ZOI) was measured in millimeters using a digital caliper. Ceftazidime (30 µg) and vancomycin (30 µg) discs were used as positive controls, while sterile water served as a negative control.

**Figure 1.**
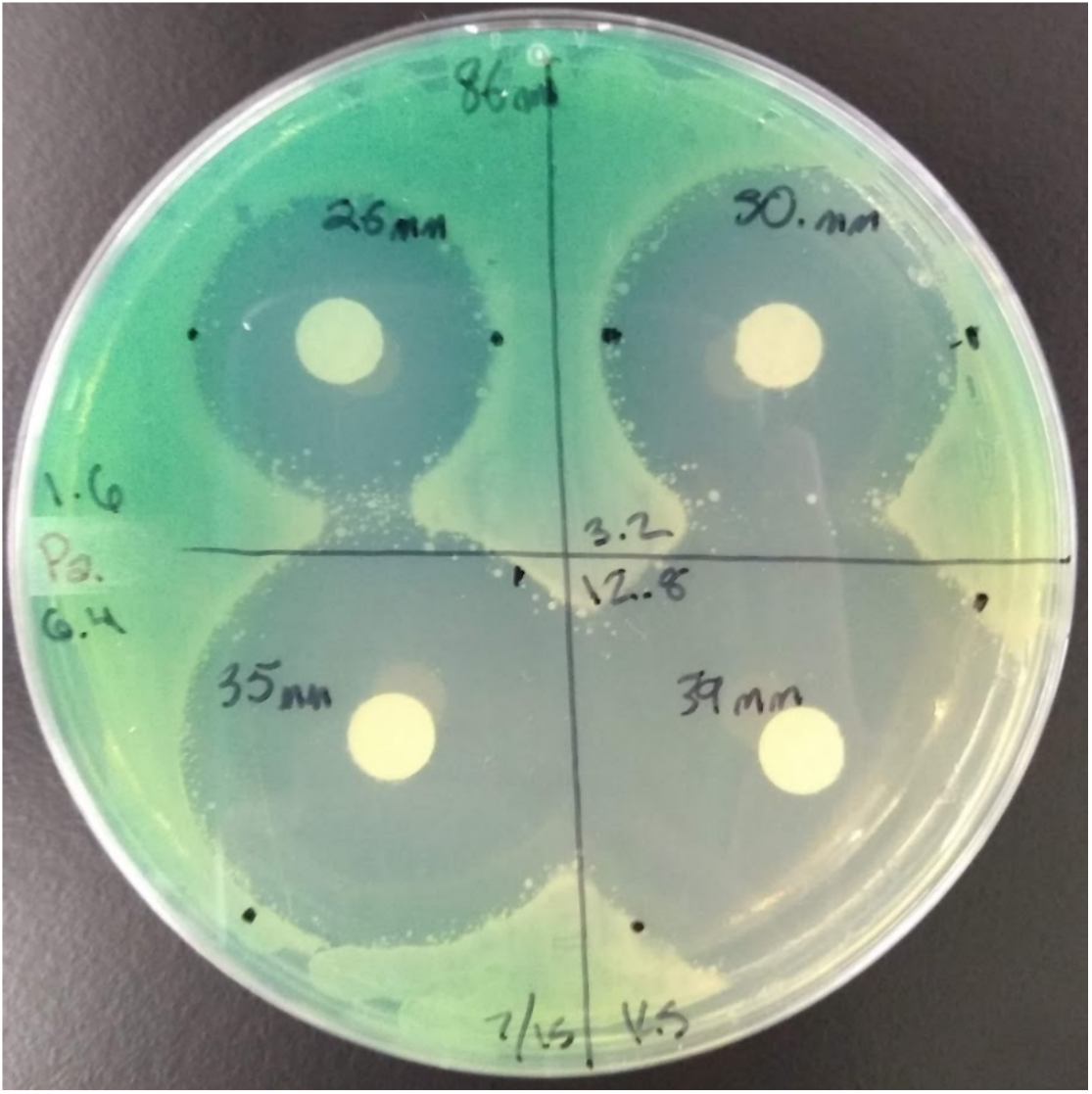
Representative disk diffusion assay demonstrating zone of inhibition. Plate results of treatment of gentamycin against *P. aeruginosa*, used here to validate assay sensitivity and confirm strain susceptibility to an aminoglycoside comparator. Gentamycin was applied at 1.6, 3.2, 6.4, and 12.8 ug per disc. Diameter of each inhibition ring was measured using digital caliper in mm.

**Figure 2.**
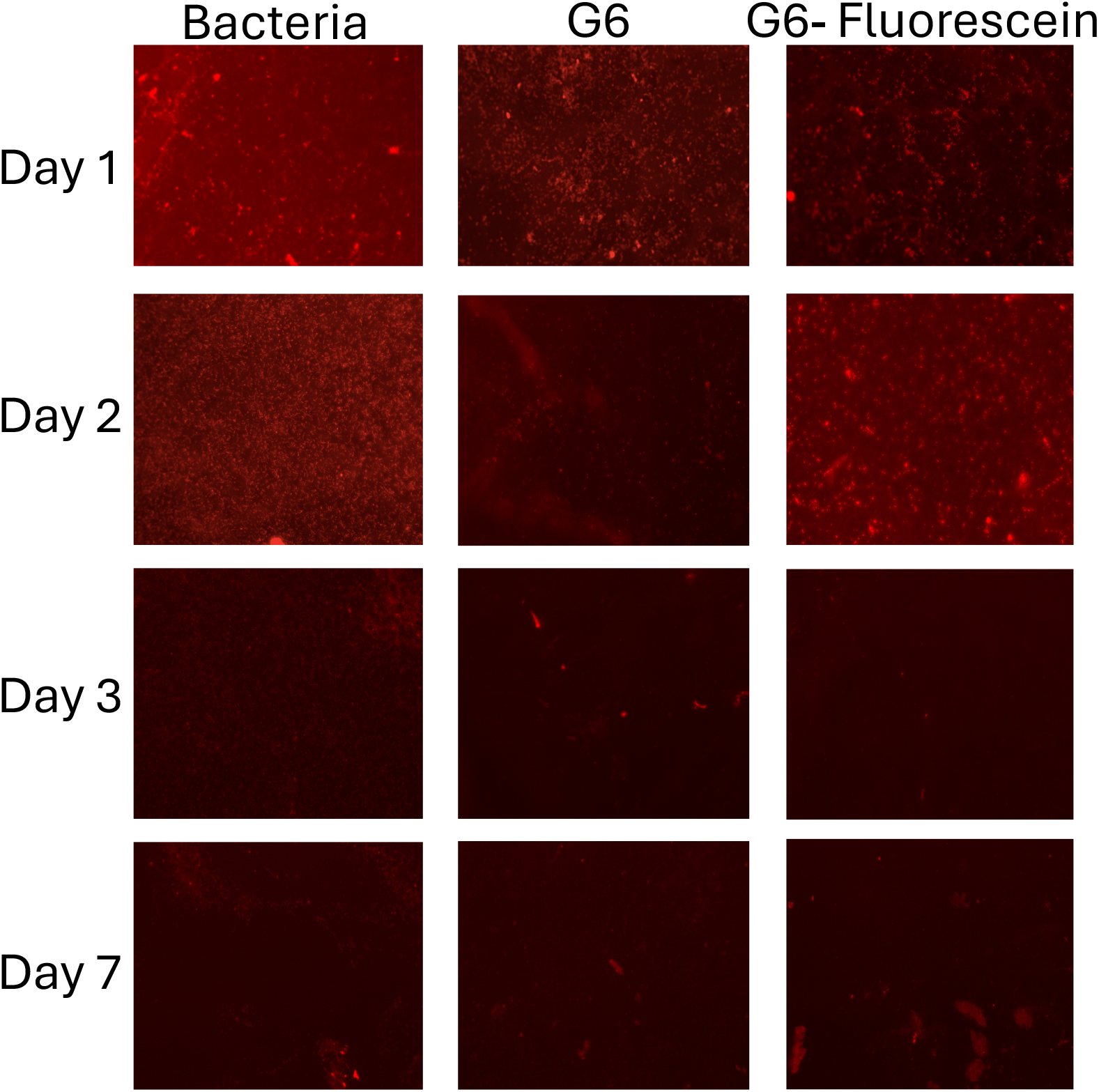
Representative photos of *P. aeruginosa* biofilm treated with dendrimers G6, G6-F (fluorescein-conjugated G6 dendrimer), or bacteria only control, and stained with SYPRO Ruby Biofilm Matrix Stain (which selectively labels extracellular matrix proteins). Images taken on days 1, 2, 3, and 7 after inoculation, each image taken under 40x objective using M5000 florescence microscope. Red florescence signifies the presence of biofilm. All images were acquired under identical exposure settings; brightness was not adjusted prior to quantification.

### Antimicrobial Susceptibility Testing

The minimum inhibitory concentration (MIC) and minimum bactericidal concentration (MBC99) of G6 and G9 were determined using the broth microdilution method following Clinical and Laboratory Standards Institute (CLSI) guidelines. Briefly, bacterial suspensions at ∼5 × 10^5^ CFU/mL were exposed to dendrimer concentrations ranging from 2 to 1024 µg/mL in 96-well plates. MIC was defined as the lowest concentration that inhibited visible bacterial growth after 24 h incubation at 37°C. To determine MBC99, aliquots from wells showing no growth were plated onto LB agar, and the lowest dendrimer concentration resulting in a 99% reduction in CFU was recorded.

### Checkerboard Synergy Assay

The synergistic effects of G6 and G9 with ceftazidime (*E. coli, P. aeruginosa*) and vancomycin (MSSA, MRSA) were assessed using a checkerboard microdilution assay. Two-fold serial dilutions of the antibiotics and dendrimers were prepared in a 96-well format, and bacterial suspensions were added to achieve a final inoculum of ∼5 × 10^5^ CFU/mL. Plates were incubated at 37°C for 24 h. The fractional inhibitory concentration index (FIC) was calculated as:

FIC = (MIC of Drug A in combo / MIC of Drug A alone) + (MIC of Drug B in combo MIC of Drug B alone)

FIC ≤ 0.5 indicated synergy, 0.5-4.0 indicated an additive effect, and ≥4.0 suggested antagonism.

### Biofilm Formation Inhibition Assay

Biofilm formation was assessed using a 96-well crystal violet assay. Briefly, *P. aeruginosa* was grown in LB or TSB at a 1:100 dilution in 96-well plates and incubated at 37°C under static conditions. To evaluate dendrimer-mediated biofilm inhibition, G6 and G9 were added at concentrations ranging from 2 to 1024 µg/mL at the time of inoculation. Plates were incubated for 1, 7, or 14 days, with media changes every 48 h. After incubation, non-adherent cells were removed, and biofilms were stained with 0.05% crystal violet for 15 min. Excess stain was washed off, and bound crystal violet was solubilized with 30% acetic acid. Absorbance was measured at 570 nm using a microplate reader.

### Biofilm Formation Inhibition Comparison Assay

To compare the dendrimers to known antimicrobial agents, LL37 and Indolicidin (Dürr et al., 2006; Falla et al., 1996), *P. aeruginosa* was seeded in 96 well plates in 5 different treatments. Plates were then incubated for 1, 2, or 6 days at 37°C under static conditions. After incubation, media was then removed and a Crystal Violet assay was preformed like what is seen above.

### Biofilm Disruption Assay

To assess the ability of dendrimers to disrupt preformed biofilms, *P. aeruginosa* biofilms were established in 96-well MBEC plates (Minimum Biofilm Eradication Concentration assay) for 1 or 7 days at 37°C. Following incubation, the lids of these MBEC plates were moved to a new plate containing dendrimer treatments (G6, G6-F) for 24 h. After treatment, biofilm biomass was quantified using the crystal violet assay as described above in the biofilm formation inhibition Assay section.

### Biofilm Inhibition Assay Using Fluorescent Imaging

To assess dendrimer interactions with bacterial biofilms, fluorescence microscopy was performed. Biofilms were grown on 1 mm glass coverslips placed in 48-well plates, sterilized, and inoculated with *P. aeruginosa* at a 1:100 dilution in TSB. Biofilms were allowed to form for 24, 48, or 72 h before treatment with G6 or G9. Post-treatment, biofilms were stained with SYPRO Ruby stain for 30 min in the dark. Coverslips were washed with sterile deionized water and mounted onto glass slides for imaging. Fluorescence images were captured in a series using an M5000 fluorescence microscope under a 40× objective. Image analysis was performed using ImageJ to quantify fluorescence intensity and dendrimer localization. Image brightness was not adjusted before quantification, and regions of interest were selected using lasso tool and artifacts with abnormal florescence present were excluded. In total, around 90% of each image was analyzed.

### Dendrimer Colocalization Experiment

*P. aeruginosa* was grown in a process like what was seen above for the fluorescent imaging experiment. The treatments used were bacteria only as a negative control, then 2x and 3x the concentration of dendrimers that had been used in the florescence experiment. Once a Biofilm had developed on the glass coverslips inside of the wells, they were removed, washed with DI water, and imaged using GFP setting of the M5000 microscope.

### Statistical Analysis

All experiments were performed in triplicate and repeated at least three independent times. Data are presented as mean ± standard deviation (SD). Statistical significance was determined using one-way ANOVA followed by Tukey’s post-hoc test for multiple comparisons using an online calculator (https://www.socscistatistics.com/tests/anova/default2.aspx). A p-value < 0.05 was considered statistically significant.

## RESULTS

### 1. Antimicrobial Susceptibility of G6 and G9 Dendrimers

The intrinsic antibacterial activity of dendrimers G6 and G9 was evaluated against *Escherichia coli, Pseudomonas aeruginosa*, methicillin-sensitive *Staphylococcus aureus* (MSSA), and methicillin-resistant *S. aureus* (MRSA). Both dendrimers exhibited minimal inhibitory capacity, with MIC values ≥1024 µg/mL for all strains tested (Supplementary Tables 1 and 2). MBC_99_ values were similarly elevated, indicating limited bactericidal activity at concentrations tested.

Disc diffusion assays further supported these findings, as no measurable zones of inhibition were observed for either dendrimer at concentrations up to 512 µg/mL. In contrast, clear inhibitory zones were produced by comparator antibiotics, such as gentamycin which produced a zone of inhibition with a diameter of 39 mm at 12.8 μg (Figure 1, Figure 3). Collectively, these data indicate that G6 and G9 possess limited intrinsic antibacterial activity under standard testing conditions.

**Figure 3.**
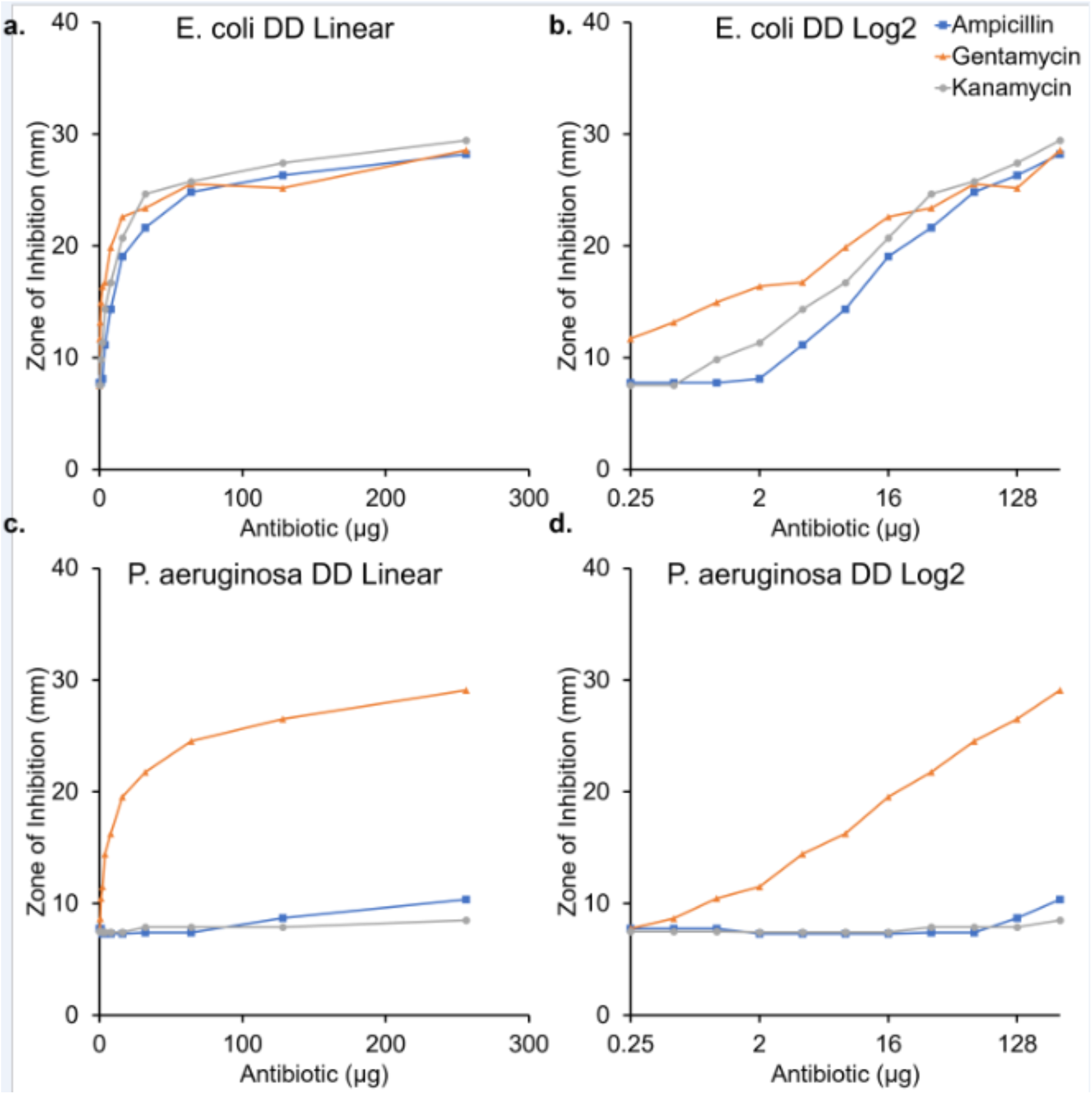
Zone of inhibition (ZOI, mm) produced by ampicillin, gentamycin, and kanamycin against *E. coli (*panels a, b) and *P. aeruginosa* (panels c, d) at increasing treatment concentrations, used to identify strain susceptibility profiles. Panels (a) and (c) both present these data plotted on a linear scale, for *E. coli* and *P. aeruginosa* respectively. Panels (b) and (d) are both presented on a log_2_ scale, for *E. coli* and *P. aeruginosa* respectively. This assay is used to establish assay validity, confirming reference strains exhibit the expected susceptibility patterns.

### 2. Synergistic Interactions Between Dendrimers and Antibiotics

#### 2.1 Synergy with Ceftazidime Against Gram-negative Species

Checkerboard assays were conducted to determine whether dendrimers enhanced susceptibility to ceftazidime in *E. coli* and *P. aeruginosa*. G9 demonstrated marked synergistic interactions, reducing the effective MIC of ceftazidime by several two-fold dilutions. *E. coli*: G9 exhibited synergistic activity with ceftazidime, with FIC indices ranging from 0.13 to 0.26 (Supplementary Table 5). The lowest FIC (0.13) corresponded to a ceftazidime concentration of 0.016 µg/mL in combination with 128 µg/mL G9. *P. aeruginosa*: G9 synergy was also observed, with FIC values between 0.31 and 0.63, indicating a reproducible reduction in ceftazidime MIC compared to antibiotic alone (Supplementary Table 5).

In contrast, G6 showed predominantly additive interactions, with FIC indices clustering near 0.5-1.0 for *E. coli* and >1.0 for *P. aeruginosa*, suggesting minimal synergy (Supplementary Table 3).

MBC_99_ checkerboard analyses confirmed these trends, with G9 lowering ceftazidime bactericidal thresholds to a greater extent than G6 (Supplementary Tables 4 and 6).

#### 2.2 Interactions with Vancomycin Against Gram-positive Species

Synergy between dendrimers and vancomycin was assessed in MSSA and MRSA. G6 and Vancomycin: FIC indices ranged from 1.0 to 4.5 across MSSA and MRSA, indicating primarily additive to antagonistic interactions (Supplementary Table 11). G9 and Vancomycin: MSSA displayed occasional additive effects (minimum FIC ≈ 1.06), while MRSA showed no synergy (Supplementary Table 13).

No combinations demonstrated FIC ≤ 0.5, indicating that dendrimer-vancomycin synergy was not observed under these conditions.

### 3. Effects of Dendrimers on Biofilm Formation

Biofilm formation by *P. aeruginosa* was quantified after 1, 2, and 6 days of growth in the presence of G6 or G9 at concentrations ranging from 2 to 1024 µg/mL. These concentrations demonstrate no significant reduction in total biofilm biomass at any time point compared with untreated controls.

Biofilm mass increased predictably with incubation time, but dendrimer treatment did not alter the biomass trajectory. These results indicate that neither G6 nor G9 substantially inhibits early biofilm establishment or longer-term biofilm accumulation under these assay conditions.

### 4. Localization of Dendrimers Within Established Biofilms

Fluorescence microscopy was used to visualize dendrimer distribution within established *P. aeruginosa* biofilms. SYPRO Ruby staining demonstrated dendrimer-associated fluorescence distributed throughout the biofilm matrix following G6 treatment. Fluorescence intensity revealed significantly higher matrix-associated signal in G6-treated biofilms compared to untreated samples. (Figure 4)

**Figure 4.**
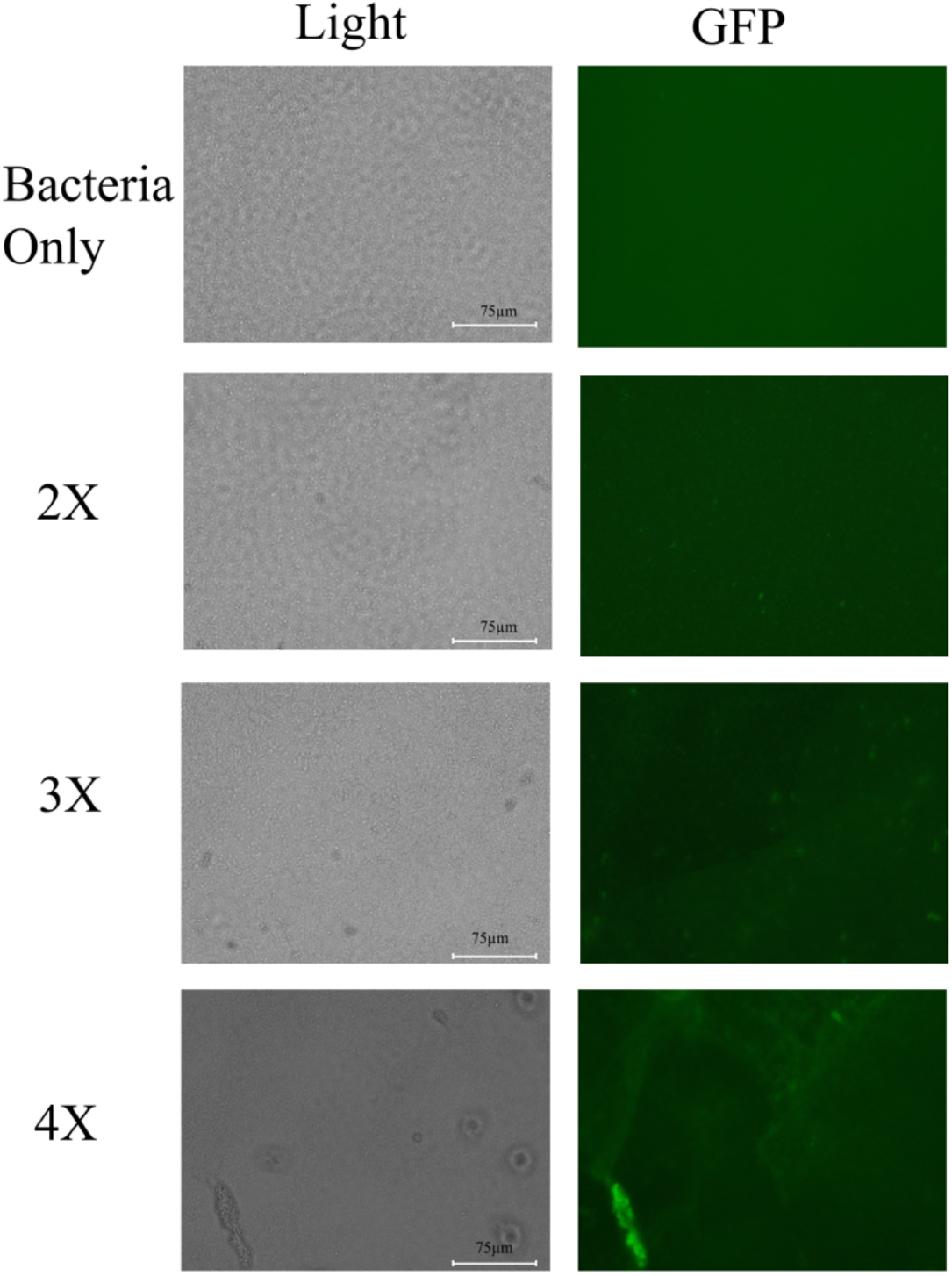
Fluorescent localization of G6-F within established *P. aeruginosa* ATCC 27853 biofilms. Biofilms were grown on glass coverslips for 48 h and then treated with G6-F, a fluorescein-conjugated G6 dendrimer, at 2×, 3×, or 4× the working concentration used in the SYPRO Ruby imaging experiment. Bacteria-only biofilms served as the negative control. Each row shows the same field imaged by light microscopy and GFP-channel fluorescence microscopy under a 40X objective using M5000 microscope. Fluorescein signal from G6-F was detected using the GFP filter set. Brighter GFP-channel signal indicates greater dendrimer-associated fluorescence in the biofilm region. The increased fluorescence at higher treatment levels is consistent with concentration-dependent G6-F accumulation within established biofilms.

Additional analyses using antimicrobial peptides (Figures 5) confirmed that matrix-associated fluorescence changes reflect differential interactions between the biofilm matrix and exogenous molecules.

**Figure 5.**
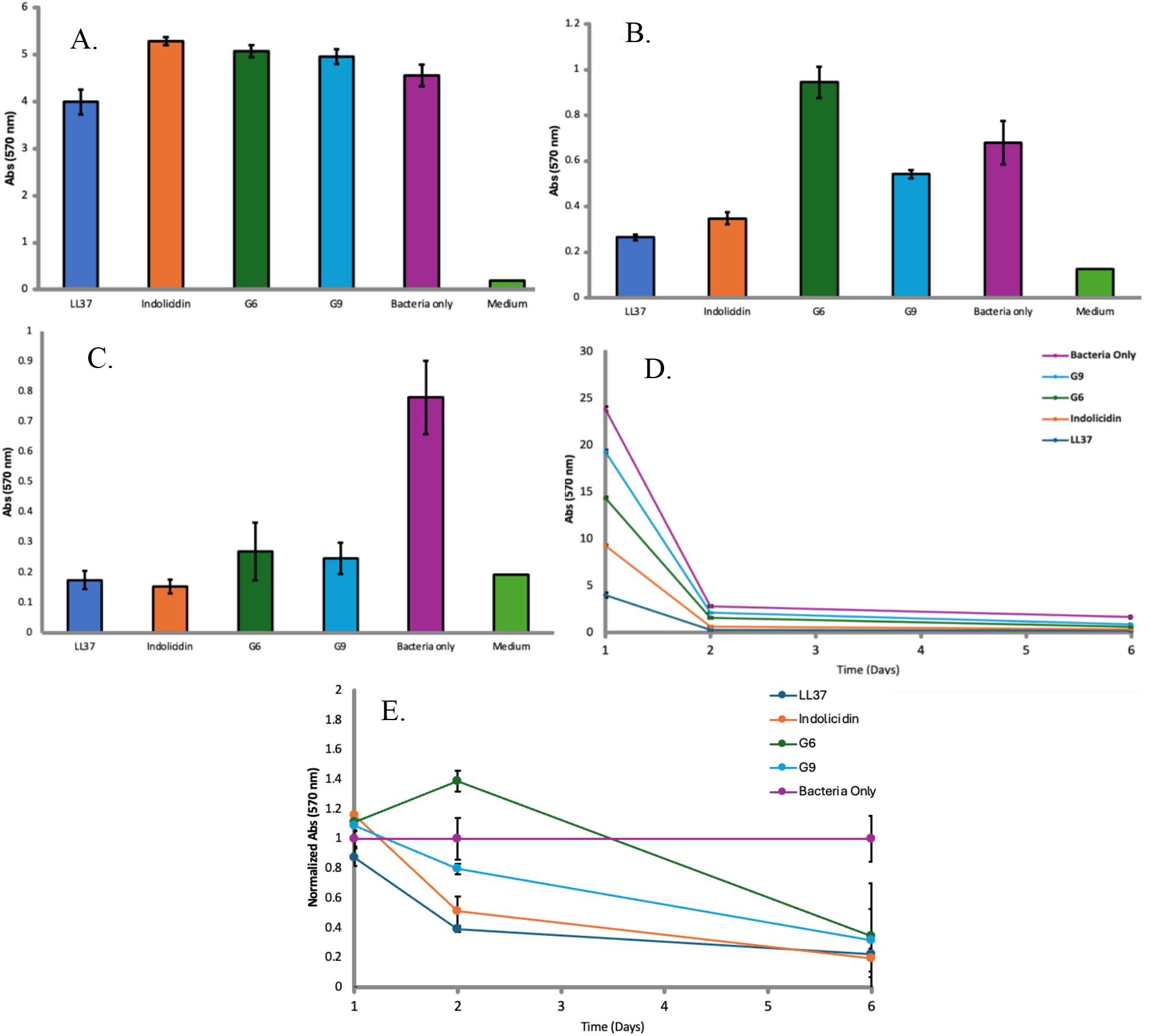
Quantification of *P. aeruginosa* biofilm biomass following treatment with dendrimers and antimicrobial peptides over 6 days of development. Biofilm biomass was measured using the crystal violet staining assay and quantified by plate reader absorbance at 570 nm after biofilm development in a 37°C sterile incubator. Treatment groups included G6 dendrimer, G9 dendrimer, LL37, Indolicidin, bacteria-only control, and media-only control where indicated. LL37 and Indolicidin were used as antimicrobial peptide comparators. Panels (a), (b), and (c) represent absorbance values from days 1, 2, and 6, respectively. Panel (d) shows all measured values plotted on the same axis, while panel (e) shows values normalized to the bacteria-only control at each corresponding time point. Error bars represent standard error for each data point. Statistical analysis by ANOVA showed significant differences among treatment groups on day 1 (p = 0.00049), day 2 (p < 0.00001), and day 6 (p < 0.00001).

### 5. Comparison of Dendrimer inhibition on Biofilm

After assaying for antibacterial properties of G6 and G9, we were interested in studying their biofilm inhibitory properties. To accomplish this, we utilized Crystal Violet Assay to measure biofilm quantity over the course of its development. *P. aeruginosa* was allowed to grow and develop biofilm in a 96 well plate for 7 days, with different treatment conditions. G6 and G9 Dendrimers were used in combination with the *P. aeruginosa*. The conditions also included positive controls Indolicidin and LL37, known antimicrobial agents. (Dürr et al., 2006; Falla et al., 1996). The negative control was the bacteria with no other treatment.

On day 1 of this experiment, the treatments were LL37, Indolicidin, G6, G9, and Bacteria only. The average absorbance on this day was 3.99 A.U., 5.29 A.U., 5.07 A.U., 4.96 A.U., and 4.56 A.U. respectively. (ANOVA p value of 0.00049) (Figure 5A). On day 2 of this experiment, the treatments were LL37, Indolicidin, G6, G9, and Bacteria only. The average absorbance on this day was 0.26 A.U., 0.35 A.U., 0.94 A.U., 0.54 A.U., and 0.68 A.U. respectively. (ANOVA p value of <0.00001) (Figure 5B). On day 6 of this experiment, the treatments were LL37, Indolicidin, G6, G9, and Bacteria only. The average absorbance on this day was 0.18 A.U., 0.15 A.U., 0.27 A.U., 0.25A.U., and 0.78 A.U. respectively. (ANOVA p value of <0.00001) (Figure 5C).

Looking at this data, the trend is therefore seen that over the course of the 6 days, many of the treatments started by causing an increase in absorption in the crystal violet assay. From there, G6 continued to increase the absorbance of the assay, while each of the other treatments decreased the absorbance compared to bacteria only. Then by the end of the experiment each of the treatments had significantly decreased the absorbance when compared to the bacteria only (Figure 5D, 5E).

### 6. Biofilm Inhibition Assay using Fluorescent Imaging

To verify these previous results, the next step was to use a SYPRO Ruby stain assay to determine growth of the biofilm with treatment of G6. Additionally, to understand the localization of these dendrimers in relation to the biofilm, G6 dendrimers with a fluorescent tag were used. SYPRO Ruby stains the matrix of biofilm, and many images were gathered in a line, then brightness data was gathered using ImageJ software.

On day 1 of this experiment, the treatments were G6 dendrimers, G6-F dendrimers, and Bacteria only. The average absorbance on this day was 64.79 A.U., 65.41 A.U., and 47.40 A.U. respectively. (ANOVA p value of 0.0057) (Figure 6). On Day 2, the treatments were again G6 dendrimers, G6-F dendrimers, and Bacteria only. The average absorbance on this day was 52.02 A.U., 35.30 A.U., and 50.99 A.U. respectively. (ANOVA p value of <0.00001) (Figure 6). On Day 3, the treatments were G6 dendrimers, G6-F dendrimers, and Bacteria only. The average absorbance on this day was 45.74 A.U., 44.85 A.U., and 53.23 A.U. respectively. (ANOVA p value of 0.014) (Figure 6). Finally, on day 7, the treatments were G6 dendrimers, G6-F dendrimers, and Bacteria only. The average absorbance by treatment group on this day was 36.19 A.U., 33.26 A.U., and 45.48 A.U. respectively. (ANOVA p value of <0.00001) (Figure 6).

**Figure 6.**
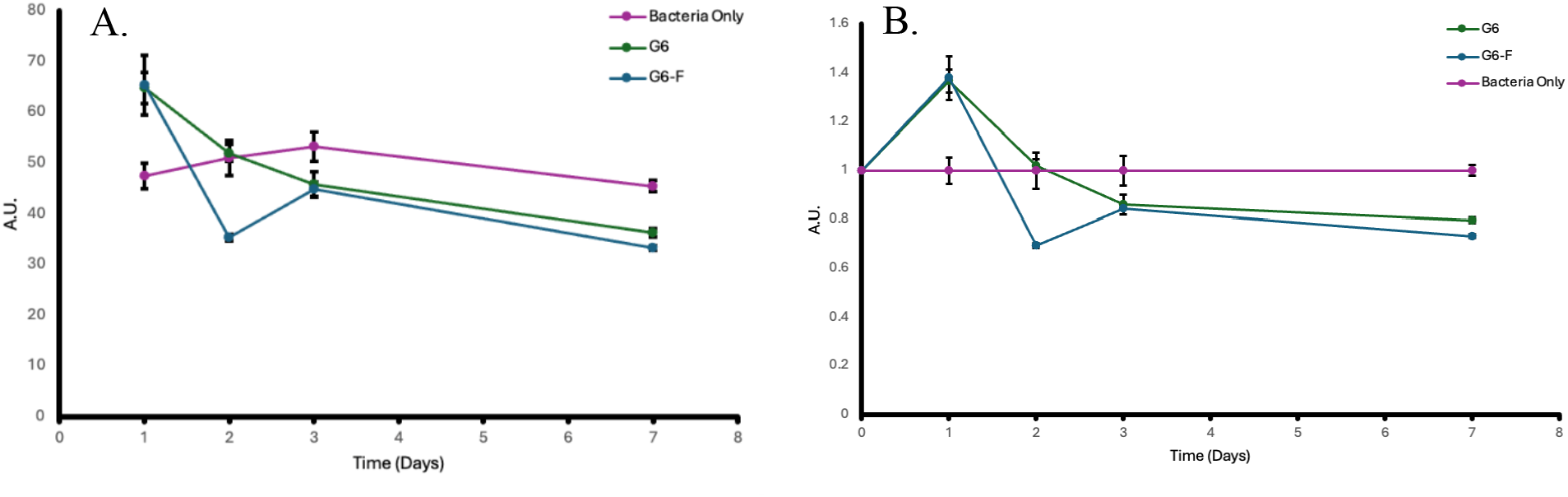
Quantification of *P. aeruginosa* biofilm matrix following G6 and G6-F treatment over 7 days of development. Biofilm matrix signal was measured using SYPRO Ruby fluorescence staining and quantified by ImageJ analysis. Treatment groups included G6 dendrimer, G6-F (fluorescein-conjugated G6 dendrimer), and bacteria-only control. Panel (A) shows raw fluorescence intensity values in arbitrary units (A.U.) across days 1, 2, 3, and 7. Panel (B) shows fluorescence values normalized to the bacteria-only control. Biofilms were developed over 7 days and imaged under consistent acquisition settings. Error bars represent standard error. Statistical analysis by ANOVA showed significant differences among treatment groups on day 1 (p = 0.0057), day 2 (p < 0.00001), day 3 (p = 0.014), and day 7 (p < 0.00001).

Overall, the trend seen here mimics the previous experiment, with dendrimer treatment initially causing biofilm increase when compared to bacteria only, while by the day 6-7 range the opposite trend is seen and there is less growth visible.

### 7. Biofilm Disruption Assay

In interest of better understanding if these dendrimers can also be applied to disrupting already formed biofilm, a biofilm disruption assay was performed by allowing a biofilm to develop over a variable number of days, then transferring that developed biofilm into a treatment of G6 and G6-F dendrimers for 1 day. A crystal violet assay was then performed on the biofilm to measure its relative quantity.

On the biofilms that were developed for 1 day before transfer, the treatments were media only, bacteria only, G6 Dendrimers, and G6-F dendrimers. The average absorbance values on this day were 0.15 Abs_470_, 0.58 Abs_470_, 0.71 Abs_470_, and 1.05 Abs_470_ respectively. (ANOVA p value of <0.00001) (Figure 7). On the biofilms that were developed for 7 days before transfer, the treatments were media only, bacteria only, G6 Dendrimers, and G6-F dendrimers. The average absorbance values on this day were, 0.15 Abs_470_, 0.94 Abs_470_, 0.87 Abs_470_, and 0.78 Abs_470_ respectively. (ANOVA p value of <0.00001) (Figure 7).

**Figure 7.**
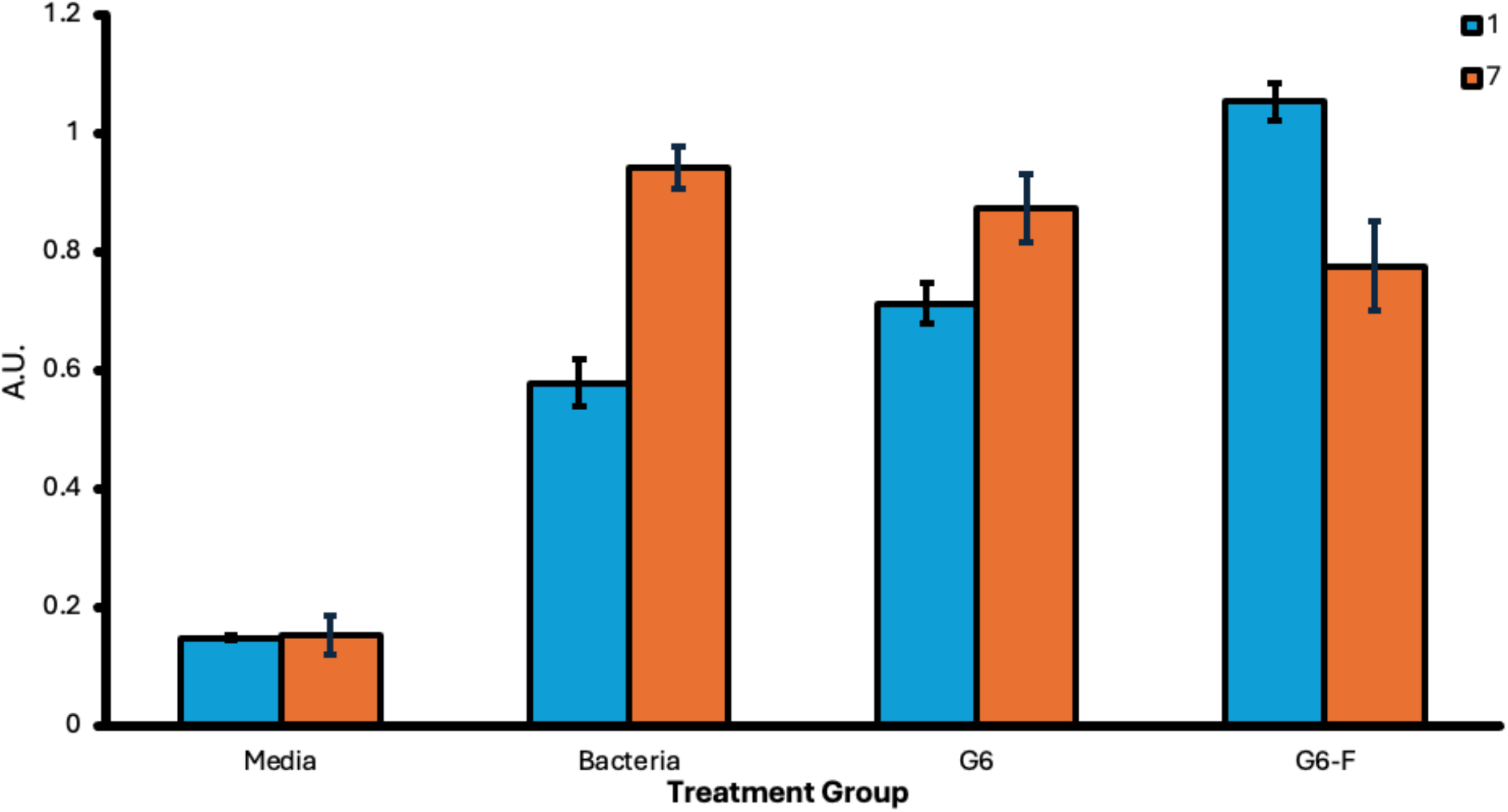
Biofilm disruption assay measuring the effect of G6 and G6-F on pre-formed *P. aeruginosa* biofilms using the crystal violet assay. Biofilms were developed in 96-well MBEC plates for either 1 day (blue bars) or 7 days (orange bars) before being transferred into new 96-well plates containing treatment. Treatment groups included media-only control, bacteria-only control, G6 dendrimer, and G6-F, a fluorescein-conjugated G6 dendrimer. After treatment, residual biofilm biomass was quantified by crystal violet staining. Higher absorbance values indicate greater remaining biofilm biomass. Error bars represent standard error. Statistical analysis by ANOVA showed significant differences among treatment groups for both 1-day biofilms (p < 0.00001) and 7-day biofilms (p < 0.00001).

The trend from this experiment shows that at longer time periods, the biofilm can be disrupted by dendrimers, while initially the dendrimers cause an increase in biofilm, comparable to what is seen above.

## DISCUSSION

This study characterized the antibacterial, synergistic, and anti-biofilm properties of two cationic dendrimers, G6 and G9, against clinically relevant Gram-negative and Gram-positive bacterial pathogens. Although both dendrimers exhibited limited intrinsic antibacterial activity, we identified important functional properties that inform their potential use as antimicrobial adjuvants rather than stand-alone therapeutics.

The observed synergy between dendrimers and ceftazidime suggests potential applications in combination therapies. The lack of direct antimicrobial activity at physiologically relevant concentrations indicates that dendrimers may be more effective as adjuvants rather than standalone agents. Their presence in biofilms with a modest but statistically significant amount of inhibition implies an alternative role in altering biofilm architecture rather than eradication. Future studies should focus on optimizing dendrimer structure to enhance biofilm penetration and antimicrobial potency.

### Limited intrinsic antibacterial activity of G6 and G9

Consistent with previous observations that dendrimer antimicrobial activity is highly dependent on surface chemistry and generation, generation number, and terminal group identity (Calabretta et al., 2007; Mintzer et al., 2012) neither G6 nor G9 produced meaningful growth inhibition of *E. coli*, MSSA, or MRSA at concentrations up to 1024 µg/mL. However, G6 and G9 dendrimers were able to significantly inhibit *P. aeruginosa* in a way that is comparable to some known antimicrobial molecules, but only after multiple days of growth. The absence of detectable zones of inhibition and the elevated MIC and MBC_99_ values (Supplementary Tables 1-2) indicate that these dendrimers do not efficiently disrupt bacterial viability under standard conditions. This behavior is not uncommon among higher-generation cationic dendrimers, which may exhibit steric shielding of surface charges due to back-folding of terminal amines, paradoxically reducing membrane interaction efficiency despite increased charge density (Tomalia et al., 2007). This limited intrinsic potency underscores the importance of evaluating dendrimers not solely as direct antimicrobials but also as molecular scaffolds capable of modulating microbial susceptibility or biofilm physiology.

### Generation-dependent synergy with ceftazidime in Gram-negative bacteria

The most notable finding of this study was the strong synergy observed between G9 and ceftazidime against *E. coli* and *P. aeruginosa*. FIC indices as low as 0.13 (Supplementary Table 5) demonstrate significant potentiation of β-lactam activity, suggesting that G9 enhances antibiotic access or disrupts membrane stability in a manner complementary to ceftazidime’s mechanism of action. Cationic macromolecules are known to displace divalent cations (Mg^2+^ and Ca^2+^) from lipopolysaccharide (LPS) binding sites in the Gram-negative outer membrane, increasing permeability and facilitating antibiotic ingress (Hancock and Chapple, 1999; Nikaido, 2003). The synergy observed here between G9 and ceftazidime is consistent with this model, wherein outer membrane destabilization by G9 lowers the effective barrier to β-lactam access to periplasmic penicillin-binding proteins. In contrast, G6 displayed primarily additive or indifferent interactions (Supplementary Tables 3-4), indicating that synergistic potential increases with dendrimer generation or charge density.

No synergy was observed between either dendrimer and vancomycin in MSSA or MRSA (Supplementary Tables 11-14). This organism-specific disparity underscores the structural and physiological differences between Gram-negative and Gram-positive envelopes; synergistic effects may require interactions with the outer membrane or periplasmic space that are absent in Gram-positive organisms. Collectively, these data suggest that dendrimer-antibiotic synergy is both generation-dependent and pathogen-specific, with the most promising applications in the treatment of Gram-negative infections.

### Dendrimers penetrate biofilms and disrupt biomass on a protracted timeframe

Although neither dendrimer significantly inhibited initial (Day 1) biofilm formation (Figures 5-6), fluorescence imaging revealed prominent dendrimer localization to established *P. aeruginosa* biofilms (Figure 4). The distribution of dendrimer-associated fluorescence throughout the matrix suggests that these molecules can penetrate biofilm structures rather than being excluded by the extracellular polymeric matrix.

Interestingly, dendrimer penetration corresponded to reductions in biomass after longer development (day 7), indicating that matrix access alone is sufficient to disrupt biofilm stability or viability in a minor way at the tested concentrations. Penetration is an essential prerequisite for any molecule intended to enhance antibiotic delivery or modulate biofilm tolerance. These findings are therefore consistent with a model in which G9, and to a lesser extent G6, may serve as biofilm-permeable carriers or sensitizers, amplifying the effects of co-administered antibiotics even when they exert little direct biofilm inhibition on their own.

### Implications for antimicrobial adjuvant development

Taken together, these results support the concept that dendrimers with limited bactericidal activity can still play a valuable role in antimicrobial therapy by functioning as adjuvants that potentiate existing antibiotics and penetrate protective biofilm matrices. The strong synergy between G9 and ceftazidime suggests potential clinical utility in treating biofilm-associated or multidrug-resistant Gram-negative infections, particularly where ceftazidime susceptibility is borderline or compromised. Additionally, dendrimer penetration into biofilms raises the possibility of improved antibiotic delivery or matrix modification, both of which warrant further mechanistic investigation.

### Study limitations

This study has several limitations. First, all experiments were conducted using reference strains; clinical isolates with diverse resistance mechanisms may exhibit different susceptibility patterns. Second, while dendrimer penetration into biofilms was visualized qualitatively and semi-quantitatively, additional high-resolution imaging and molecular analyses are needed to determine how dendrimers interact with specific matrix components. Third, synergy studies were limited to two antibiotics; expanding to other β-lactams, aminoglycosides, or polymyxins may reveal broader adjuvant potential. Finally, cytotoxicity and pharmacokinetic properties of these dendrimers were not assessed here and must be evaluated before translational application.

### Conclusions and future directions

G6 and G9 dendrimers exhibit some minimal direct antimicrobial activity but show generation-dependent synergy with ceftazidime in Gram-negative pathogens and penetrate mature biofilms while disrupting biomass. This study demonstrates that cationic dendrimers G6 and G9 possess limited intrinsic antibacterial activity but exhibit properties that support their potential use as antimicrobial adjuvants. Although neither dendrimer eradicated bacterial pathogens at physiologically relevant concentrations, G9 displayed strong synergy with ceftazidime in *E. coli* and *P. aeruginosa*, markedly enhancing antibiotic potency. Both dendrimers also colocalized with established *P. aeruginosa* biofilms, indicating that they can access protective extracellular matrices even without reducing total biomass. These findings position higher-generation cationic dendrimers, particularly G9, as promising candidates for further development as antimicrobial adjuvants and support a therapeutic model in which dendrimers may augment the activity of conventional antibiotics and enhance their performance in biofilm-associated infections. Future work should focus on mechanistic studies of dendrimer-antibiotic interactions, evaluation against resistant clinical and multidrug-resistant isolates, and structural optimization to enhance antimicrobial potency while preserving biofilm penetration capabilities, which will be essential to advancing dendrimers as viable components of combination antimicrobial therapy.

## Acknowledgements

We thank the Discovery Institute, Seattle Washington, and the Roth family for their kind support of this research. We also thank Dr. Hyun Ho Chung for his generous donation of the G6 and G9 Dendrimers for this study.

